# The Genetic History of Northern Europe

**DOI:** 10.1101/113241

**Authors:** Alissa Mittnik, Chuan-Chao Wang, Saskia Pfrengle, Mantas Daubaras, Gunita Zarina, Fredrik Hallgren, Raili Allmäe, Valery Khartanovich, Vyacheslav Moiseyev, Anja Furtwängler, Aida Andrades Valtueña, Michal Feldman, Christos Economou, Markku Oinonen, Andrejs Vasks, Mari Tõrv, Oleg Balanovsky, David Reich, Rimantas Jankauskas, Wolfgang Haak, Stephan Schiffels, Johannes Krause

**Affiliations:** Max Planck Institute for the Science of Human History, Jena, Germany; Institute for Archaeological Sciences, Archaeo- and Palaeogenetics, University of Tübingen, Tubingen, Germany; Department of Archaeology, Lithuanian Institute of History, Vilnius; Institute of Latvian History, University of Latvia, Riga, Latvia; The Cultural Heritage Foundation, Västerås, Sweden; Archaeological Research Collection, Tallinn University, Tallinn, Estonia; Peter the Great Museum of Anthropology and Ethnography (Kunstkamera) RAS, St. Petersburg, Russia; Archaeological Research Laboratory, Stockholm University, Stockholm, Sweden; Finnish Museum of Natural History - LUOMUS, University of Helsinki, Finland; Independent researcher; Research Centre for Medical Genetics, Moscow, Russia; Vavilov Institute for General Genetics, Moscow, Russia; Department of Genetics, Harvard Medical School, Boston, Massachusetts 02115, USA; Broad Institute of Harvard and MIT, Cambridge, Massachusetts 02142, USA; Howard Hughes Medical Institute, Harvard Medical School, Boston, Massachusetts 02115, USA; Department of Anatomy, Histology and Anthropology, Vilnius University, Vilnius, Lithuania; School of Biological Sciences, The University of Adelaide, Adelaide SA-5005, South Australia, Australia

## Abstract

Recent ancient DNA studies have revealed that the genetic history of modern Europeans was shaped by a series of migration and admixture events between deeply diverged groups. While these events are well described in Central and Southern Europe, genetic evidence from Northern Europe surrounding the Baltic Sea is still sparse. Here we report genome-wide DNA data from 24 ancient North Europeans ranging from ∼7,500 to 200 calBCE spanning the transition from a hunter-gatherer to an agricultural lifestyle, as well as the adoption of bronze metallurgy. We show that Scandinavia was settled after the retreat of the glacial ice sheets from a southern and a northern route, and that the first Scandinavian Neolithic farmers derive their ancestry from Anatolia 1000 years earlier than previously demonstrated. The range of Western European Mesolithic hunter-gatherers extended to the east of the Baltic Sea, where these populations persisted without gene-flow from Central European farmers until around 2,900 calBCE when the arrival of steppe pastoralists introduced a major shift in economy and established wide-reaching networks of contact within the Corded Ware Complex.

---

Recent genetic studies of ancient human genomes have revealed a complex population history of modern Europeans involving at least three major prehistoric migrations^1–6^ that were influenced by climatic conditions and availability of resources as well as the spread of technological and cultural innovations and possibly diseases^7,8^. However, how and when they affected the populations of the very north of the European continent surrounding today's Baltic Sea, where the archeological record shows a distinct history to that of Central and Southern Europe, has yet to be comprehensively studied on a genomic level.

The archeological record of the eastern Baltic and Scandinavia shows that settlement by mobile foragers started only after retreat of the glacial ice sheets around 11,000 years before present^9^. To the west and south, hunter-gatherers (Western Hunter-Gatherers or WHG) sharing a common genetic signature already occupied wide ranges of Europe from Iberia to Hungary for several millennia^1,2,5,10,11^. They were shown to be descended from foragers appearing in Europe after around 14,000 years ago in a population turnover coinciding with the warming period of the Bølling-Allerød interstadial, possibly emerging from a southern refugium they inhabited since the Glacial Maximum and replacing preceding foraging populations^5,12^. From further to the east in the territory of today's Russia, remains of Mesolithic foragers (Eastern Hunter-gatherers or EHG) have been studied^2,4^. They derived the majority of their ancestry, referred to as Ancient North Eurasian ancestry (ANE), from a population related to the Upper Paleolithic Mal'ta boy found in Siberia (MA1)^6,13^. Late Mesolithic foragers excavated in central Sweden, called Scandinavian Hunter-Gatherers (SHG)^1,2^, were modeled as admixed between WHG and EHG^6^. Foraging groups along the eastern Baltic coast increasingly relied on marine resources during the 8^th^ and 7^th^ millennium calBCE and lived in more permanent settlements than their surrounding contemporaries^14,15^.

The following Early Neolithic period, starting around 6,000 calibrated radiocarbon years before Common Era (calBCE), saw the transition from foraging to a sedentary agricultural lifestyle with the expansion of farmers out of Anatolia following the Danube and Mediterranean coast into Central and Southern Europe where they existed in parallel and admixed with local foragers for the following two millennia^1,4,6,16,17^. This development reached South Scandinavia at around 4,000 calBCE with the farmers of the Funnel Beaker Culture (TRB; from German *Trichterbecher*) who gradually introduced cultivation of cereals and cattle rearing. At the transition to the northern Middle Neolithic, around 3,300 calBCE, an intensification of agriculture is seen in Denmark and western Central Sweden accompanied by the erection of megaliths and changes in pottery and lithic technology, while settlements in eastern Central Sweden increasingly concentrated along the coast and economy shifted toward marine resources such as fish and seal. A gradual change in material culture can be seen in the archeological assemblages of these coastal hunter-gatherers, known as the Pitted Ware Culture (PWC) with early pottery resembling the Funnel beakers in shape. Analysis of ancient genomes from PWC and megalithic Middle Neolithic TRB context in Central Sweden has shown that PWC individuals retain the genetic signature of Mesolithic hunter-gatherers while the TRB farmers’ ancestry can mainly be traced back to Central European farmers, albeit with substantial admixture from European hunter-gatherers^18–20^

The production and use of pottery, in Central and southern Europe often seen as part of the ‘Neolithic package’, was already common among foragers in Scandinavia during the preceding Mesolithic Ertebølle phase. Similarly in the eastern Baltic, where foraging continued to be the main form of subsistence until at least 4,000 calBCE^15^, ceramics technology was adopted before agriculture. Recent genome wide data of hunter-gatherers from the Baltic Narva Culture revealed genetic continuity with the preceding Mesolithic inhabitants of the same region as well as influence from the more northern EHG^21^.

The Late Neolithic is seen as a major transformative period in European prehistory, accompanied by changes in burial customs, technology and mode of subsistence as well as the creation of new cross-continental networks of contact seen in the emergence of the pan-European Corded Ware Complex (CWC, ca. 2,900 to 2,300 calBCE) in Central^2^ and northeastern Europe^21^. Studies of ancient genomes have shown that CWC were genetically closely related to the pastoralist Yamnaya Culture from the Pontic-Caspian steppe, bringing with them a genetic component that was not present in Europe previously^2,3^. Genomes from the CWC of Central Germany suggest that this new genetic component replaced around 75% of the local Middle Neolithic genetic substrate^2^. Presumably this ‘steppe’ genetic component spread in the subsequent millennia of the Final Neolithic and Bronze Age throughout Europe and can be seen in today's European populations in a decreasing northeast to southwest gradient. Intriguingly, modern eastern Baltics carry the most WHG ancestry of all Europeans^1^, supporting the theory of a remnant Mesolithic hunter-gatherer population in this region that left a lasting genetic impact on subsequent populations^22^.

Here we genetically investigate these dynamics of population turnover and continuity at the northern fringe of prehistoric European occupation. In the most comprehensive ancient DNA study in Northern Europe to date, we present novel genome-wide data from 24 ancient individuals spanning 7,000 years of prehistory from the Baltics, Russia and Sweden. We show that the settlement of Scandinavia by hunter-gatherers likely took place via two different routes, and that the first introduction of farming was brought about by migration of Central European farmers around 4,000 calBCE. In the eastern Baltics, foraging remained the dominant economy, corresponding with a genetic continuity of the population up until around 3,000 calBCE, when we see the first major shift towards agro-pastoralism brought on by migrations from the Pontic steppe as opposed to Central Europe.

## Results and Discussion

### Samples and archaeological background

The skeletal remains studied here were recovered from 25 archaeological sites in the territory of modern Lithuania, Latvia, Estonia, Karelia (Russia) and Sweden dating from around 7,500 to 200 calBCE (Fig.1, Supplementary Information Section1, Supplementary Data Table 1). In total, we analysed DNA from 105 human remains. Twenty-four samples that exhibited good DNA preservation were selected for deeper shotgun sequencing or SNP capture (Extended Data Table 1). In the latter case, we enriched samples for a panel of around 1.24 million single nucleotide polymorphisms (SNPs) via in-solution capture^4,23^ to an average coverage of 0.04 to 8.8-fold on targeted SNPs and co-analysed with previously published data from over 300 ancient and around 3800 modern individuals^6^.

**Figure 1.**
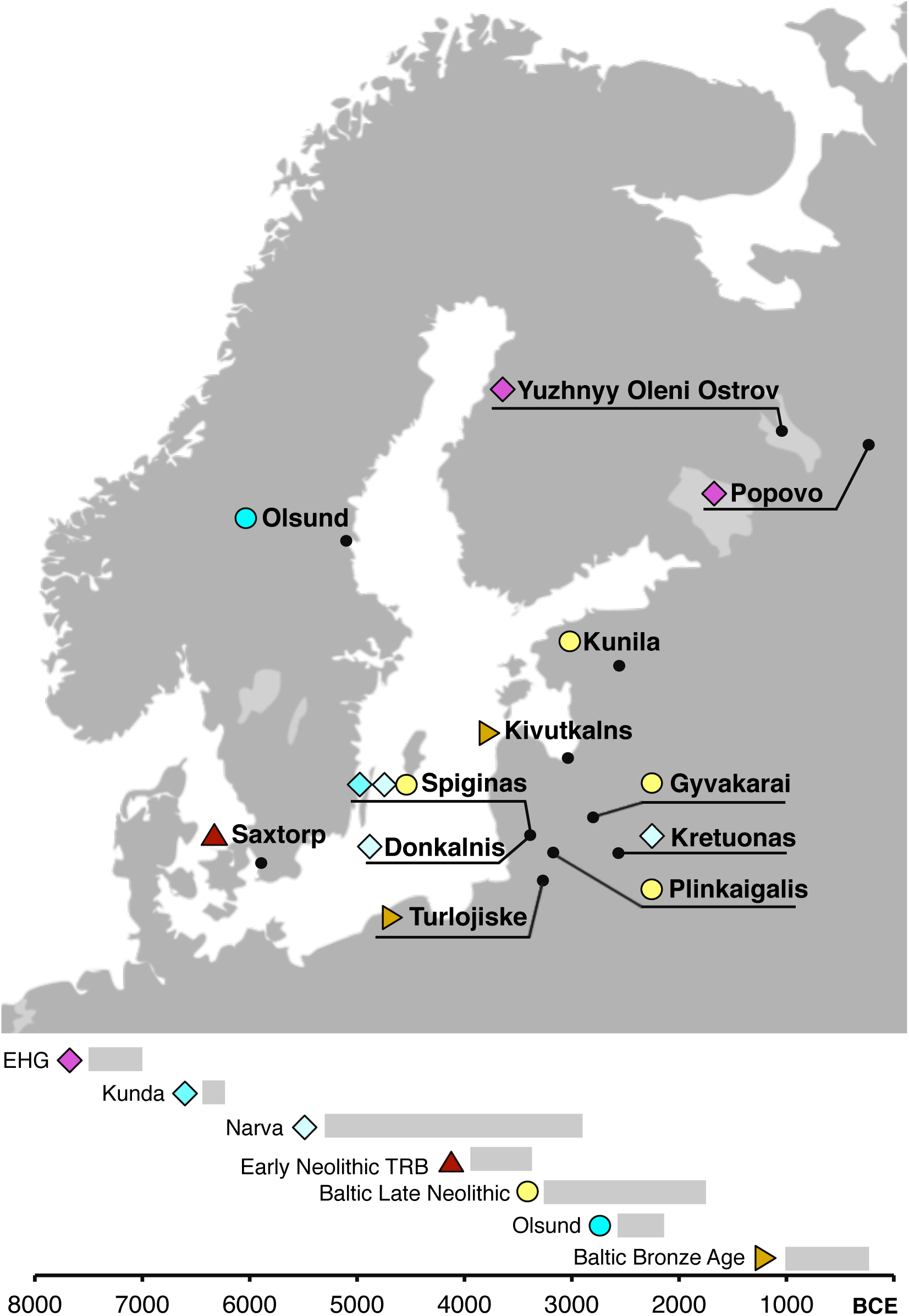
Sampling locations and dating of ancient Northern European genomes introduced in this study. Chronology based on calibrated radiocarbon dates or contextual dating, see Supplementary Information Section 1.

The 24 final samples fall into seven broad groups: First, two Mesolithic hunter-gatherers from northwestern Russia (ca. 7,500 to 5,500 calBCE), adding to the EHG genomes from this context that are already available^4^; secondly, a Mesolithic hunter-gatherer from Lithuania (6440–6230 calBCE) associated with the Kunda Culture, whose archeological assemblages found in Lithuania, Latvia, Estonia and adjacent parts of Russia. Four individuals (ca. 4,700 to 4,200 calBCE) were assigned to the Narva Culture of eastern Baltic ceramist hunter-gatherers. Five individuals from Lithuania and Estonia were dated to the Late Neolithic (ca. 3200–1750 calBCE) and grouped together with the already published Estonian genome^3^ as Baltic Late Neolithic (LN). Nine samples from Latvia and Lithuania were attributed to the Baltic Bronze Age (Baltic BA) and date to ca. 1000–200 calBCE. From southern Sweden we analyse two farmers (ca. 3900–3400 calBCE) from the Early Neolithic TRB Culture (EN TRB), the earliest agricultural population in Scandinavia. As previously genetically analysed TRB individuals date to a period one millennium after the Neolithisation in Southern Scandinavia, these samples address the open question whether the first introduction of farming around 4,000 BCE was driven by newcomers or by local groups involving later gene-flow from Central European farmers. One sample from northern Sweden (Olsund, ca. 2570–2140 calBCE) dates to the Latfige Neolithic but was found without associated archeological assemblages.

### Affinities of northern Mesolithic hunter-gatherers

To gain an overview of the broad genetic affinities of our samples, we projected all 24 ancient genome-wide datasets onto principal components, constructed from 1007 modern individuals from a diverse set of West Eurasian contemporary populations. We also projected previously published ancient samples^4^ onto this plot and investigated the relationship of ancient Baltics to other ancient populations in Europe. The two samples from Karelia cluster with previously published Mesolithic EHG (Fig. 2A), exhibit similar composition of their genetic makeup (Fig. 2b) and share most genetic drift since divergence from Africa with EHG as shown by outgroup f3-statistics (Extended Data Figures 1 and 2). We therefore grouped them together with EHG for all following analyses. Model-based clustering using ADMIXTURE also shows that EHG carry a genetic component (green component in Fig. 2b) that is maximized in hunter-gatherers from the Caucasus (CHG) and shared with Neolithic farmers from Iran and Steppe populations from the Bronze Age, suggesting some common ancestry for these populations, consistent with previous results^21^.

**Figure 2.**
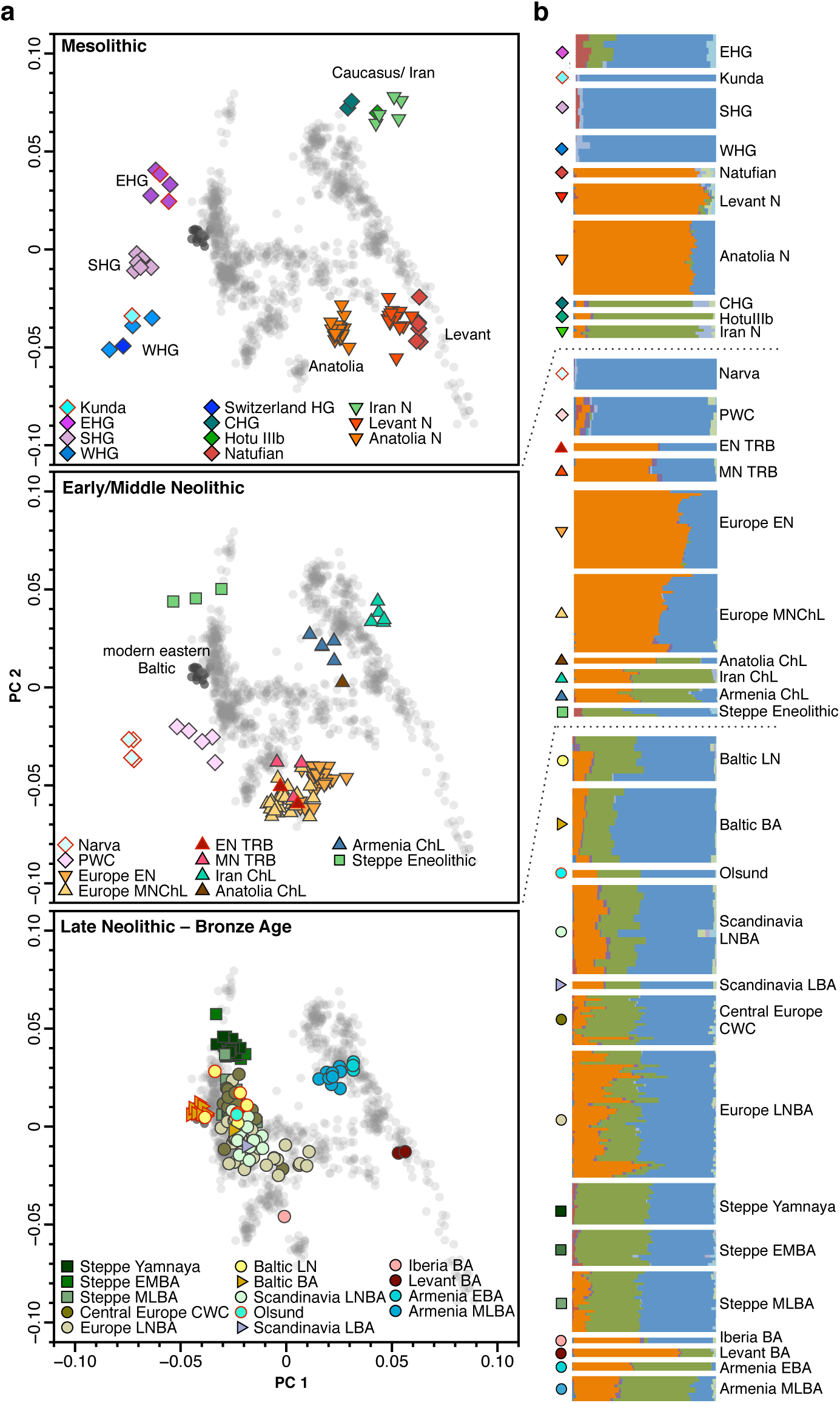
PCA and ADMIXTURE analysis reflecting three time periods in Northern European prehistory. **(a)** Principal components analysis of 1007 present-day West Eurasians (grey points, modern Baltic populations in dark grey) with 282 projected published ancient and 24 ancient north European samples introduced in this study (marked with a red outline). Population labels of modern West Eurasians are given in Supplementary Information Fig. 2 and a zoomed-in version of the European Late Neolithic and Bronze Age samples is provided in Extended Data Fig. 4. (b) Ancestral components in ancient individuals estimated by ADMIXTURE (k=11).

Despite its geographically vicinity to EHG, the eastern Baltic individual associated with the Mesolithic Kunda culture shows a very close affinity to WHG in all our analyses, with a small but significant contribution from EHG or SHG, as revealed by significant D-statistics of the form *D*(Kunda, WHG; *EHG/SHG,* Mbuti) (Z>3; Supplementary Information Table S2).

SHG appear intermediate between WHG/Kunda and EHG in PCA space, and the statistic *D*(SHG,WHG; EHG, Mbuti) is strongly significant for excess allele sharing of SHG and EHG (Z=12.8). Neither the Kunda individual nor SHG exhibit the major ADMIXTURE component shared between EHG and CHG (green in Figure 2b), bringing into question a direct contribution of EHG into the Mesolithic individuals from Scandinavia and the eastern Baltic. However, using *qpWave* we cannot reject the previously published result of SHG being formed by admixture of WHG and EHG^6^ (58±2% WHG with 42±2% EHG; p=0.55; Supplementary Information Table S7). Both EHG and SHG share a non-negligible component in ADMIXTURE analysis that is maximized in some modern Native American populations which points towards ancient North Eurasian ancestry, as represented by the MA1 and AG3 samples from Palaeolithic Siberia^13^ (crimson component in Fig. 2a). Indeed, D-statistics show that EHG and SHG share significantly more alleles with MA1 and AG3 than both Baltic and Western HG (Supplementary Information Table S9). Additionally, mtDNA haplogroups found among EHG point toward an eastern influence: R1b in UzOO77 was previously found in the Palaeolithic Siberian AG3^5^ and a haplogroup related to branches within the C1 clade, which appears today in highest frequencies in northeast Asia and the Americas, was described in several samples from Yuzhnyy Oleni Ostrov^24,25^. Furthermore, in SHG the derived variant of the EDAR allele was discovered, which is found today in high frequency in East Asians and Native Americans^4^.

In contrast to EHG and SHG, Kunda can be modelled as directly derived from WHG (p=0.18) (Supplementary Information Table S6). The almost complete absence of the additional ancestry shared by SHG and EHG to the south of the Baltic Sea suggests that it was brought into Scandinavia via a northern route through Finland and admixed in Scandinavia with a WHG-like population that derived from a migration northward over the land-bridge that connected Denmark and southern Sweden at the time, a scenario that is in concordance with the archeological record^26^.

### An eastern Baltic refugium of European hunter-gatherers

The results for the Kunda individual are mirrored in the four later eastern Baltic Neolithic hunter-gatherers of the Narva culture (Fig. 2) and further supported by the lack of significantly positive results for the D-statistic *D*(Narva, Kunda; *X,* Mbuti) (Supplementary Information Table S2) demonstrating population continuity at the transition from Mesolithic to Neolithic, which in the eastern Baltic region is signified by a change in networks of contacts and the use of pottery rather than a stark shift in economy as seen in Central and Southern Europe^15^. That Narva individuals derive directly from Kunda without additional admixture cannot be rejected (p=0.12), however it can also be accounted for by admixture of Kunda with either EHG, SHG or WHG (Supplementary Information Table S4). Specifically we see a greater proportion of possible admixture into the two samples excavated at the more eastern site Kretuonas (13±3% EHG or 33±7% SHG) than into the two Narva individuals from the more western sites (3±3% EHG or 7±6% SHG) (Supplementary Information Table S5). This substructure within the eastern Baltic foragers might have been present already in the Mesolithic, and evades our detection as we only have one sample from this time period.

Similar to the other Mesolithic hunter-gatherers, all our Baltic foragers carry the derived *HERC2* allele which codes for light iris color, and like SHG and EHG they already possess an increased frequency of the derived alleles for SLC45A2 and SLC24A5, coding for lighter skin color (Extended Data Table 2). The two male Narva individuals carry Y-chromosomal haplogroups of the I2a1 clade (Supplementary Information Section 3, Extended Data Table 1) and Y-haplogroup I has been most commonly found among WHG and SHG^1,5^.

The Narva individual Spiginas1 (dated to ca. 4440–4240 cal BCE) belongs to a mitochondrial haplogroup of the H branch providing the first direct evidence that this branch was present among European foragers without gene-flow from farmers (Extended Data Table 1). Notably, in addition to haplogroup H, the maternal lineages seen in eastern Baltic samples (n=31; Extended Data Figure 5) encompass all of the major haplogroups identified in complete mtDNA genomes from Holocene Scandinavian and western European hunter-gatherers (n=21:U2, U5a, U5b)^12^, as well as haplogroup U4 which has been found in high frequency in Mesolithic foragers from Russia^24^ and K1, a derivate of the U8 branch found in Scandinavian foragers^20^, suggesting a large census size of Baltic hunter-gatherers to maintain such genetic diversity. We see in Baltic foragers no genomic evidence of gene-flow from Central European farmers or any Y-chromosomal or mitochondrial haplogroups that are typical for them, suggesting that any traces of agriculture and animal husbandry in the Baltic Early and Middle Neolithic were due to local development or cultural diffusion^15^.

### Early Neolithic farmer migration into Scandinavia

In contrast to the Eastern Baltic, we see clear evidence of a movement of agriculturalists into Southern Sweden already around 4,000 calBCE. The individuals associated with the Early Neolithic TRB culture (EN TRB) cluster with Middle and Early Neolithic farmers from Europe on the PCA (Fig. 2A) and in the ADMIXTURE analysis exhibit the component maximized in Levantine early farmers (orange component in Figure 2b). The statistic *D*(EN_TRB, Middle Neolithic Central Europe; *X*, Mbuti) does not yield significantly positive results and EN TRB can be modeled as derived from a single source identical with Middle Neolithic (MN) Central Europe (p=0.26; Supplementary Information Table S6). Models involving a linear combination of MN Central Europe and different groups of northern European hunter-gatherers are not rejected, but the admixture proportions estimated for the latter do not significantly deviate from zero (e.g. 4±6% for SHG at p=0.26; Supplementary Information Table S7). Previous studies^2,27^ have shown a resurgence of WHG ancestry during the European Middle Neolithic coinciding with the expansion of farming to regions previously inhabited by hunter-gatherers. We demonstrate here that early Swedish farmers, similarly, derive from a mix between Early Farmers and WHG, without additional gene-flow from foraging groups represented by SHG from Central Sweden. However, as no forager samples from South Sweden, Denmark or Northern Germany have been studied as of yet we cannot say if the genetic substrate was more WHG-like in these regions and if local admixture might have played a greater role in the TRB culture. Nevertheless, our results suggest that the first introduction of agriculture into South Sweden was mainly facilitated by demic diffusion from Central Europe.

The previously published succeeding farmers of the Middle Neolithic (MN) TRB culture in West Sweden^19^ appear as directly descended from the EN TRB, with no significant positive results for *D*(MN_TRB, EN_TRB; *X,* Mbuti) and *qpWave* analysis using only EN TRB as source for MN TRB are not rejected (p=0.86).

The PWC individuals, who were contemporaneous to the MN TRB but relied mainly on marine resources, appear intermediate between SHG and Middle Neolithic farming cultures on the PCA (Fig. 2A) and the only models not rejected with *qpWave* involve a two-way admixture between SHG and EN TRB (91±3% SHG and 9±4% EN TRB Supplementary Information Table S7). By this model, PWC is largely genetically continuous to SHG, which is congruent with their similarities in economy but somewhat at odds with archaeological models indicating continuity between EN TRB and the later PWC^28^. We suggest that the continuity between EN TRB and the later PWC seen in archeological assemblages could be due to contact between foraging and farming groups with gene-flow from the latter into the former. Admixture in the other direction from Scandinavian foraging groups into early farmers cannot be excluded in our analyses but is not significantly different from zero (Supplementary Information Table S7).

### New networks of contact during the Late Neolithic and Bronze Age

The substantial population movement at the beginning of the third millennium calBCE, during the European Late Neolithic and Early Bronze Age (LNBA), which affected the genetic makeup of Eastern and Central Europe and Scandinavia^2,3^, made its mark in the eastern Baltic region as well, as seen in our five samples from Lithuania and Estonia dated to this period and the previously published individual from Sope, Estonia^3^. All Baltic Late Neolithic (LN) individuals (ca. 3,200 to 1,750 calBCE) fall in PCA space in the diffuse European LNBA cluster formed by individuals admixed between Early and Middle Bronze Age (EMBA) pastoralists from the Yamnaya culture of the eastern Pontic Steppe and Middle Neolithic European farmers (Fig. 2A) and carry the genetic component that was introduced into Europe with this pastoralist migration (green in Fig. 2B). This impact is also reflected in the uniparental markers where we see novel mitochondrial haplogroups (I, J, T2, W), not found in the preceding foragers, in half of our samples (Extended Data Figure 5), and I2a Y-chomosomal haplogroups replaced by R1a types (Supplementary Information Section 3, Extended Data Table 1).

*qpWave* estimates that the Baltic LN samples, when analysed as a population, are consistent with being derived from the same source as Central European CWC samples (p=0.12; Supplementary Information Table S4) and no significant positive hits appear for the statistic *D*(Baltic_LN, CordedWare_Central; X, Mbuti). Analysed individually, however, this model is rejected for three LN samples: Gyvakarai1 and Plinkaigalis242, which is dated to the very beginning of the LN, are instead consistent with being derived from the same source as EMBA Steppe pastoralists (p=0.41 and p=0.19, respectively; Supplementary Information Table S4), which corresponds with their ADMIXTURE profiles that lack the early farmer component also missing in EMBA Steppe samples (orange component in Fig. 2b). Coinciding with this steppe-like genetic influx is the first evidence of animal husbandry in the eastern Baltic^15^, suggesting import of this technology by an incoming steppe-like pastoralist population independent of the agricultural societies that were already established to the south and west.

Furthermore, the individual Spiginas2, which is dated to the very end of the Late Neolithic, has a higher proportion of the hunter-gatherer ancestry, as seen in ADMIXTURE (darker blue component in Fig. 2b), and is estimated to be admixed between 78±4% Central European CWC and 22±4% Narva (Supplementary Information Table S6). A reliance on marine resources persisted especially in the north-eastern Baltic region until the end of the Late Neolithic^29^ and in combination with the proposed large population size for Baltic hunter-gatherers a ‘resurgence’ of hunter-gatherer ancestry in the local population through admixture between foraging and farming groups is likely, and has been described for the European Middle Neolithic^2,30^.

We observe that the increased affinity to Baltic hunter-gatherers continues in the more recent samples from the Baltic BA (dated between ca. 1,000 and 230 calBCE), that cluster together on the PCA in the same space occupied by modern Lithuanians and Estonians, shifted from other Europeans to WHG and Baltic hunter-gatherers (Fig. 2A). The statistic *D*(Baltic_BA, Baltic_LN; Narva, Mbuti) is strongly significant (Z=11.7; Supplementary Information Table S2) demonstrating the increase in allele sharing with local hunter-gatherers in the Baltic populations after the Late Neolithic. Additionally, D-statistics provided significantly positive results for *D*(Baltic_BA, Baltic_LN; *X*, Mbuti) when *X* was replaced by various agricultural populations of Europe and the Near East (Supplementary Information Table S2) which suggests that the formation of the Baltic BA gene pool was not completed by admixture between the Baltic CWC population and foragers but involved additional gene-flow from outside the Baltic territory. Direct evidence of this exists in the Baltic BA sample that appears as an outlier on the PCA, falling within the larger European LNBA cluster instead of the tight Baltic BA cluster (Fig. 2a) and archaeological evidence supports that the site Kivutkalns, which is represented by eight of our individuals, was a large bronze-working center located on a trade route that opened to the Baltic Sea on the west and led inland following the Daugava river^31^. The Baltic BA was furthermore the first eastern Baltic population to show an increased frequency of the derived *LCT* allele, which is responsible for lactase persistence, i.e. the ability to digest unprocessed dairy (Extended Data Table 2). This rise in frequency could be due to either gene-flow carrying the allele into the region or a strong positive selection for this phenotype^4^.

The individual from Olsund in north-eastern Sweden was dated to the Late Neolithic (ca. 2,600 to 2,100 calBCE) when agriculture had been introduced to the coastal areas of Northern Sweden with the Battle Axe Culture, the regional variant of the CWC, while foraging persisted as an important form of subsistence. The remains were found without any associated artifacts, but in close proximity to a site where the assemblage showed a mix between local hunter-gatherer traditions and CWC influence^32^.

On the PCA this sample falls within the European LNBA cluster (Fig. 2A) and similarly to other individuals from this cluster displays the three components derived from WHG, CHG and Neolithic Levant (Fig. 2b) providing genomic evidence for the influence of both the early Neolithic and LNBA expansions having reached as far as northern Sweden in the third millennium.

*qpWave* rejects both Scandinavia LNBA and Baltic LN as single sources for Olsund and all tested models of two-way admixture between Baltic LN or Scandinavia LNBA and other contemporaneous groups (Supplementary Information Table S5). However, in all tests Baltic LN consistently provides higher p-values over Scandinavia LNBA suggesting the former population in more closely related to the potential source of the Olsund individual. This could indicate that the route of CWC expansion into Northern Sweden might have not been northward from Southern Scandinavia but instead westward across the Baltic Sea either by boat or over the frozen sea during winter^33^. Assemblages similar to the early CWC of Sweden have been found in south-western Finland, across the Bothnian Sea^34,35^ which could be considered a geographically closer source than Southern Scandinavia.

### Gene-flow into the eastern Baltic after the Bronze Age

Despite the close clustering of modern eastern Baltic populations with Baltic BA on the PCA plot and Lithuanians and Estonians exhibiting the highest allele sharing for ancient Baltic populations with any modern population (Extended Data Figure 2), Baltic BA as a single source for either modern Lithuanians or Estonians is rejected (Supplementary Information Table S4). The statistic *D*(Lithuanian, Baltic_BA; X, Mbuti) reveals significant positive results for many modern Near Eastern and Southern European populations which can be caused by Lithuanians having received more genetic input from populations with higher farmer ancestry after the Bronze Age (Supplementary Information Table S8). As this applies to nearly all modern populations besides Estonians, especially for Central and Western Europe, limited gene-flow from more south-western neighbouring regions is sufficient to explain this pattern.

In contrast, the statistic *D*(Estonian, BA_Baltic; *X,* Mbuti) gives the most significant positive hits for East Asian and Siberian populations (Supplementary Information Table S8) as previously suggested^2^. This might be connected to the introduction of the Y-chromosomal haplogroup N that in Europe is found in highest frequencies in Finland and the eastern Baltic states, and in similar high frequencies in the Uralic speaking populations of the Volga-Ural region^36^. The spread of N into north-eastern Europe was proposed to have happened with speakers of Uralic languages from the east who contributed to the male gene pool of eastern Baltic populations and left linguistic descendants in the Finno-Ugric languages Finnish and Estonian^37,38^. As we do not see Y-haplogroup N in any of the male samples from Lithuania and Latvia dated as late as 230 calBCE we propose that this element was brought into the gene pool of the more southern region of the Baltic coast after the Late Bronze Age.

### Conclusion

With our analyses we support the pattern seen in the archeological record of continuity between the Mesolithic and Early Neolithic hunter-gatherer populations in the territory of modern Lithuania who appear genetically similar to Western hunter-gatherers. In contrast, contemporaneous hunter-gatherers from the more northern Latvia and Estonia were closer to Eastern hunter-gatherers^21,39^. Networks of contact between the Baltic Sea and the river Volga could explain similarities seen in this region in pottery styles of hunter-gatherer groups although morphologically analogous ceramics could also have developed independently due to similar functionality^40^.

The situation appears differently in Scandinavia where a transition from foraging to agriculture in the Early Neolithic is carried by a demic diffusion from the south. Both the eastern Baltic and Scandinavia saw persistence of foraging throughout the Middle and Late Neolithic in populations that genetically largely descended from hunter-gatherer ancestors, as the resources of the Baltic Sea region, exploited through fishing and hunting, provided a beneficial environment for these groups and made it possible for them to maintain large population sizes without relying on crop cultivation.

We see a population movement into the regions surrounding the Baltic Sea with the Corded Ware Complex in the Late Neolithic that introduced animal husbandry to the eastern Baltic regions but did not completely replace local foraging societies. The presence of ancestry from the Pontic Steppe among Baltic CWC individuals without the Anatolian farming component must be due to a direct migration of steppe pastoralists that did not pick up this ancestry in Central Europe. This could lend support to a linguistic model that sees a branching of Balto-Slavic from a Proto-Indo-European homeland in the west Eurasian steppe^41–43^. As farming ancestry however is found in later eastern Baltic individuals it is likely that considerable individual mobility and a network of contact throughout the range of the CWC facilitated its spread eastward, possibly through exogamous marriage practices^44^. Conversely, the appearance of mitochondrial haplogroup U4 in the Central European Late Neolithic after millennia of absence^27^ could indicate female gene-flow from the eastern Baltic region, where this haplogroup was present at high frequency.

Potential future directions of research are the identification of the proximate ancient populations that contributed ancestry into Scandinavian and Karelian hunter-gatherers and Native Americans, and the timing of the arrival of Y-haplogroup N and with it possibly some of the Finno-Ugric languages spoken today in the eastern Baltic region and Fennoscandia.

### Online Methods

#### Screening of samples

DNA was extracted^45^ from a total of 80 ancient samples (teeth and bones) from the eastern Baltic region, ranging from the Mesolithic Kunda culture to the Late Bronze Age. From the Scandinavia (Sweden) we sampled 21 human remains from Mesolithic and early Funnel beaker (TRB) contexts. The samples and their archeological context are described in Supplementary Information Section 1 and presented in a tabular overview with sequencing results in Supplementary Data Table 1. Detailed screening methods are described in Supplementary Information Section 2.

#### Nuclear capture and sequencing

Twenty-three samples (including two previously studied Karelian samples^24^) were chosen for nuclear capture, the enrichment of ∼1.2 million SNPs informative about population ancestry. Additional uracil-DNA-glycosylase (UDG) treated libraries^46^ were prepared out of the DNA extracts of these samples. These libraries were then barcoded with sample specific index sequence combinations^47^, subsequently amplified with Herculase II Fusion (Agilent) and enriched for a targeted set of ∼1.2 million nuclear SNPs (1240k SNP set)^2,4,23^.

Enriched libraries from the eastern Baltic and Swedish samples were paired-end sequenced on a NextSeq500 at the department of Medical Genetics at the University of Tübingen using 2×150 bp reads and a HiSeq4000 at the IKMB in Kiel, Germany, using 2×150bp reads, the UDG-treated library of UzOO77 was processed at the Max Planck Institute for the Science of Human History in Jena, Germany, and was sequenced there on a HiSeq4000 for 2×76 cycles, and the UDG-treated library for Popovo2 was processed at Harvard Medical School, Boston, USA, and sequenced here on a NextSeq500 for 2×75 cycles.

Additionally, the non-UDG-treated screening library of Gyvakarai1 was paired-end sequenced on two lanes of a HiSeq4000 for 2×100 cycles, and on a full run of a NextSeq500 for 2×75 cycles. The screening library for Kunila2 was paired-end sequenced deeper on 80% of one lane of a HiSeq4000 for 2×100 cycles. Additionally, 40 μL of DNA extract of Kunila2 was converted into a UDG-treated library, and pair-end sequenced on one lane of a HiSeq4000 for 2×75 cycles. The three UDG-half libraries for Olsund were single-end sequenced on a HiSeq4000 for 77bp.

Resulting sequence data was demultiplexed and then further processed using EAGER^48^. This included mapping with BWA (v0.6.1)^49^ against UCSC genome browser's human genome reference GRCh37/hg19, and removing duplicate reads with same orientation and start and end positions. To avoid an excess of remaining C-to-T and G-to-A transitions at the ends of the reads, two bases of the ends of each read were clipped.

For each of the targeted SNP positions, the genotype was called by choosing a read at random to represent this position using the in-house genotype caller *pileupcaller.*

#### Quality control

One sample that was covered at less than 10,000 SNPs of the 1240k SNP set was excluded from further analyses (Turlojiske4).

Contamination in male samples was estimated to be under 5% by testing for heterozygosity of the X chromosome, using the software ANGSD^50^(Supplementary Data 1). For the female samples and the male sample Popovo2 that had too few reads on the X chromosome to produce a contamination estimate with ANGSD, we restricted to reads showing post-mortem damage characteristically seen in ancient DNA with the software *pmdtools*^51^ and called genotypes from these datasets. We devised a test *f_4_*(*Test,* Chimp; Sample_PMDfiltered, Sample_nonPMDfiltered) where *Test* was iterated through the modern populations Mbuti, Yoruba, Kalash, French, Karitiana, Papuan and Atayal to see if any of the samples shared more alleles with a modern population in the non-filtered data compared to the filtered, putatively authentic data, indicating contamination related to the modern population. None of the unfiltered samples showed a significant (|Z|>3) affinity to any of the modern populations.

Analyses on Saxtorp5158 were still restricted to only reads showing post-mortem damage as this sample had an elevated contamination rate on the mtDNA. Furthermore, the software *lcmlkin*^52^ estimated a coefficient of relatedness of around 0.5 between this sample and Saxtorp5164, consistent with first-degree relatives, and Saxtorp5158 was therefore excluded from population genetic analyses other than PCA to avoid bias resulting from a defined population consisting of closely related individuals.

#### Sex assignment

Genetic sex was assigned by calculating the ratio of X chromosomal and Y chromosomal coverage to autosomal coverage at the targeted SNPs (X and Y rate, respectively) as described previously^5^. Samples with an X rate between 0.7-1 and Y rate between 0-0.1 were assigned female and those with an X rate between 0.3-0.55 and Y rate between 0.4-0.7 were assigned male demonstrating the informative value of the Y rate over the X rate using this method (Supplementary Information Figure S1).

#### Population genetic analyses

Reference datasets for ancient populations are taken from the publicly available dataset used in Lazaridis et al. 2016^6^ (which includes genotypes from samples published earlier in Skoglund et al. 2014^19^, Haak et al. 2015^2^, Allentoft et al 2015^3^, Mathieson et al. 2015^4^ and Jones et al. 2015^53^, among others) and Fu et al. 2016^5^, as well as genotyping data from worldwide modern populations (Human Origins or HO dataset) published in the same publications and provided by the David Reich lab^6^. When analyzing only ancient samples we make use of the 1,196,358 SNPs targeted by the 1240k SNP capture, using the genotypes of deep shotgun sequenced modern Mbuti as the outgroup. For analyses involving modern populations we restrict to the intersection of 597,503 SNPs between the 1240k SNP set and the HO dataset).

Principle component analysis (PCA) was performed with *smartpca* in the EIGENSOFT package^54^ by constructing the eigenvectors from modern west Eurasian populations (Supplementary Information Figure S2) and projecting ancient individuals on these eigenvectors (Figure 2a, Extended Data Figure 4).

Admixture analysis (Figure 2b) was carried out with ADMIXTURE on 3769 modern and 366 ancient individuals for ancestral populations k=2 to k=20. The SNP dataset was pruned for linkage disequilibrium with PLINK using the parameters --indep-pairwise 200 25 0.5. We considered the cross-validation (CV) error and report in Figure 2b the results of k=11 where the CV error levels out at a minimum.

To quantify population affinities and admixtures suggested in the PCA and ADMIXTURE analysis, we carried out *f*-statistics using the programs *qp3Pop* and *qpDstat* in the ADMIXTOOLS suite (https://github.com/DReichLab) for *f*_3_- and *f*_4_- statistics, respectively. *f*_3_-statistics of the form *f*_3_(X,Y; Outgroup) measure the amount of shared genetic drift of population X and X after their divergence from an outgroup. Admixture *f*_3_-statistics of the form *f*_3_(Test;X,Y) indicate when significantly negative that population *Test* is intermediate in allele frequencies between populations A and B and could be considered an admixed population. This test was performed with parameter *inbreeding:YES* and can not be done for populations with less than two individuals. *f*_4_-statistics of the form *f*_4_(X,Y; *Test,* Outgroup) show if population *Test* is symmetrically related to X and Y or shares an excess of alleles with either of the two. Results are only reported for statistics based on more than 10,000 SNPs. Key *f*_4_ analyses were repeated using only transversion sites to control for the effect of deamination (Supplementary Information Tables S10 and S11).

To formally test for the number of source populations and the proportion of ancestry these contributed to our studied populations we used the *qpWave* and *qpAdm* programs from ADMIXTOOLS. The methodology of using regression of *f*_4_-statistics of a *Reference* population with various outgroups to relate its ancestry to a *Test* population is described in detail in the Supplementary Information of Haak et al.^2^ and Lazaridis et al.^6^. With *qpWave* we identified potential source populations for our population under study by testing if a set of *Left* populations (the *Test* population under study and its potential proximate source *Reference* populations) is consistent with being descended from *n* waves of admixture which have unique relationships to the *Right* outgroup populations (we used as modern outgroups Mbuti, Naxi, Han, Onge, Papuan, Lahu, Ami, Dinka, She, Atayal, Yi, Miao, and as ancient outgroups Switzerland_HG, MA1, Ust_Ishim and Kostenki14). This is given when rank *n-1* cannot be rejected (p>0.05), and rejected (i.e. more that *n* waves of admixture are needed to explain the ancestry of *Test* and *Reference*) if rank *n-1* can be rejected (p<0.05).

To estimate admixture proportions, we used *qpAdm* to model the Test population as a mixture of various source populations postulated from the *qpWave* test, setting as *Left* populations the *Test* and source populations and the various outgroups named above as the *Right* populations.

## Acknowledgments

We thank Isil Kucukkalipci, Guido Brandt, Nadin Rohland and Swapan Mallick for technical support in DNA analyses and Herve Bocherens, Vesa Palonen, Anne-Maija Forss and Igor Shevchuk for arranging sample treatment and technical support for radiocarbon analyses. We thank Henny Piezonka, Cosimo Posth and Iosif Lazaridis for helpful discussion and suggestions.

## Author contributions

A.M., R.J. and J.K. conceived the idea for the study. M.D., G.Z., F.H., R.A., V.M., V.K., O.B. and R.J. assembled skeletal material. A.M., S.P., A.F., A.A.V., M.F., C.E., M.O., D.R. and J.K. performed or supervised wet lab work. A.M., C.W. and S.P. analyzed data. AM., C.W., M.D., G.Z., F.H., M.T., R.A., M.O., W.H., S.S. and J.K. wrote the manuscript and supplements.

---

**Extended Data Figure 1.**
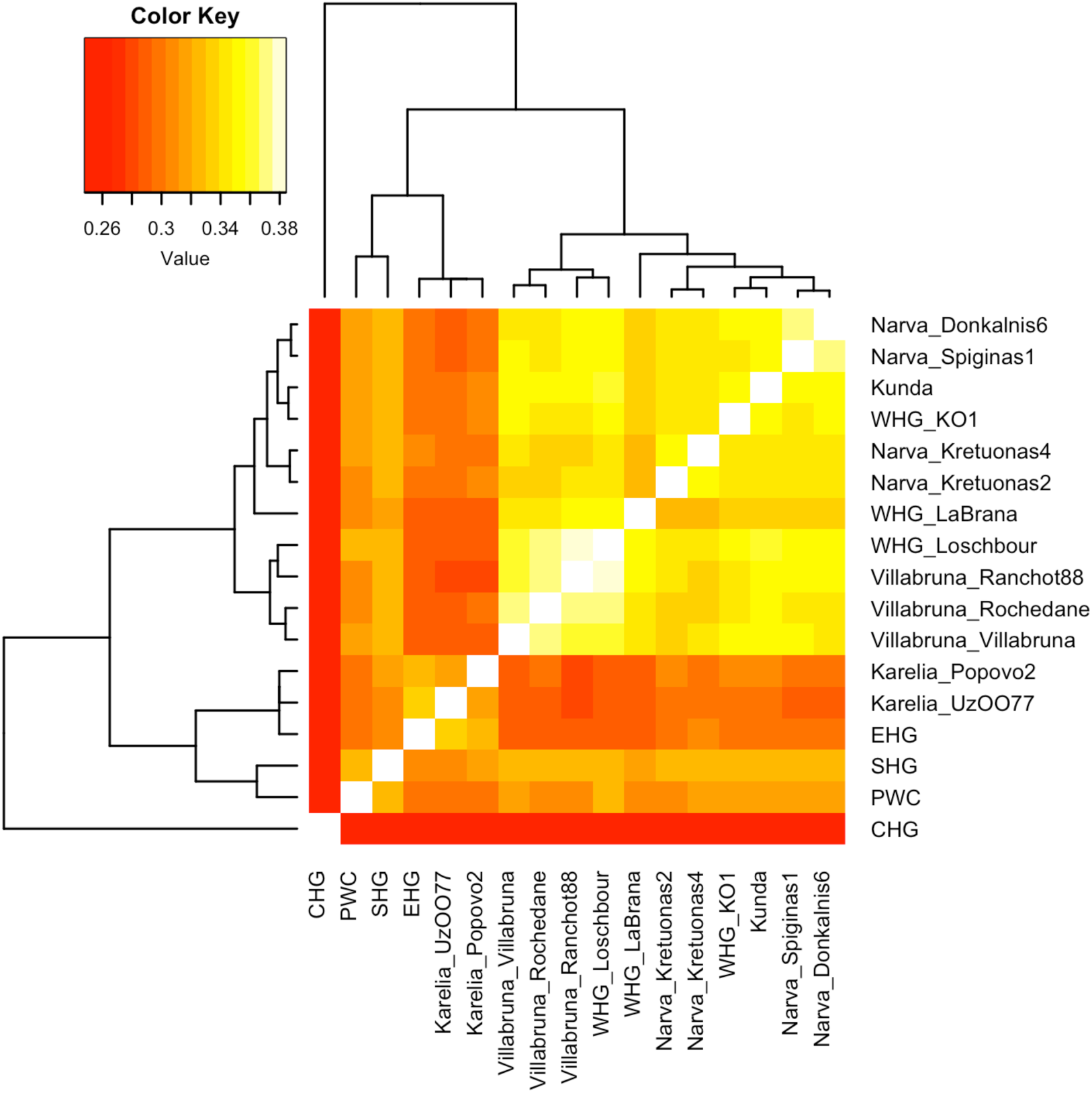
Pairwise outgroup *f*_3_-statistics of late Western Eurasian hunter-gatherers. Eastern Baltic hunter-gatherers fall within the ‘Villabruna’ cluster (including WHG) of European hunter-gatherers dating after ca. 14,000 BP. PWC and SHG cluster together and share more genetic drift with ‘Villabruna’ hunter-gatherers that with EHG.

**Extended Data Figure 2.**
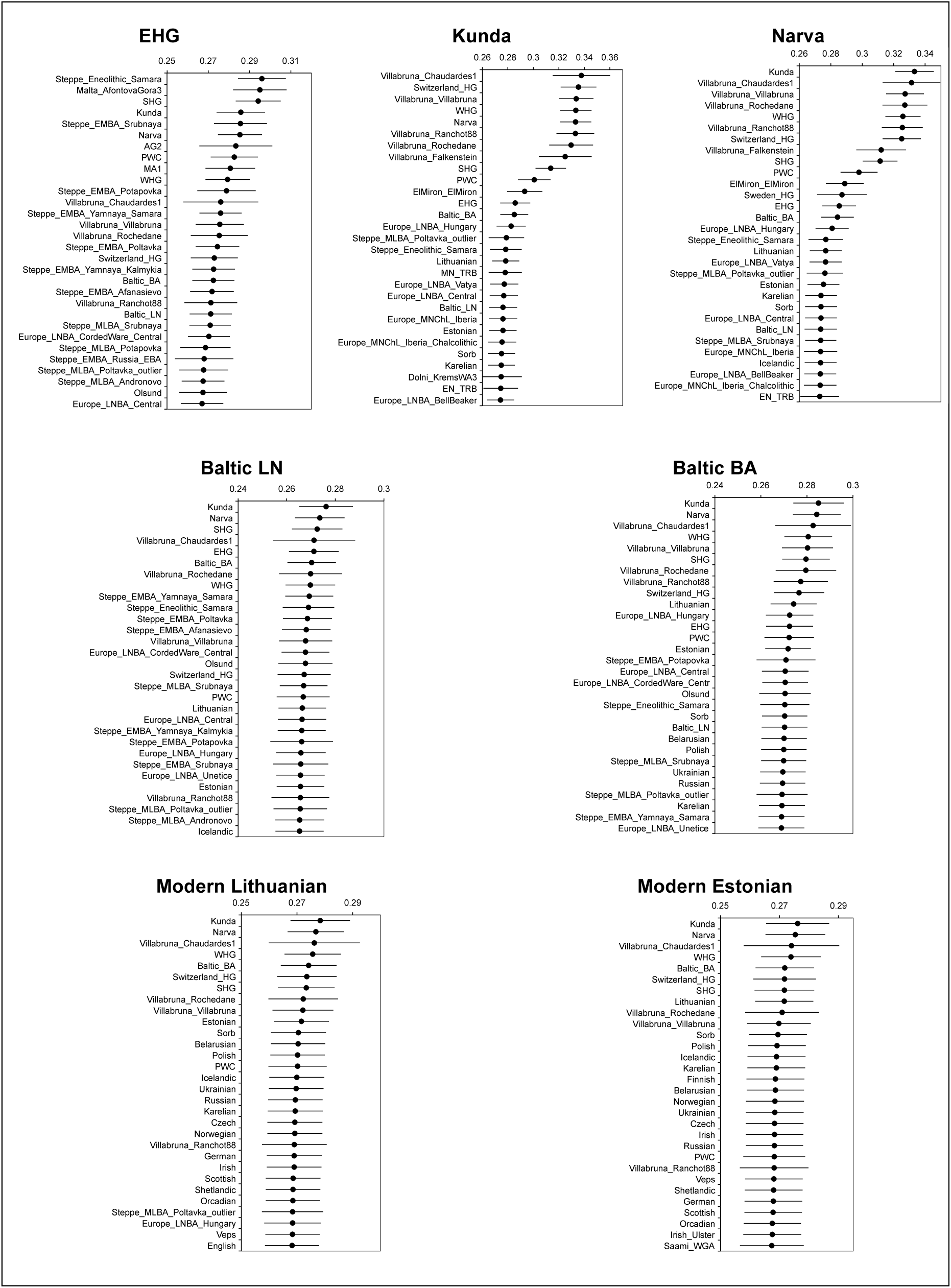
Outgroup *f*_3_-statistics for ancient and modern eastern Baltic populations. Values shown for the statistic *f*_3_(Eastern Baltic Population, *X*; Mbuti), where *X* is a modern or ancient population. The thirty highest hits are shown and error bars represent three standard errors.

**Extended Data Figure 3.**
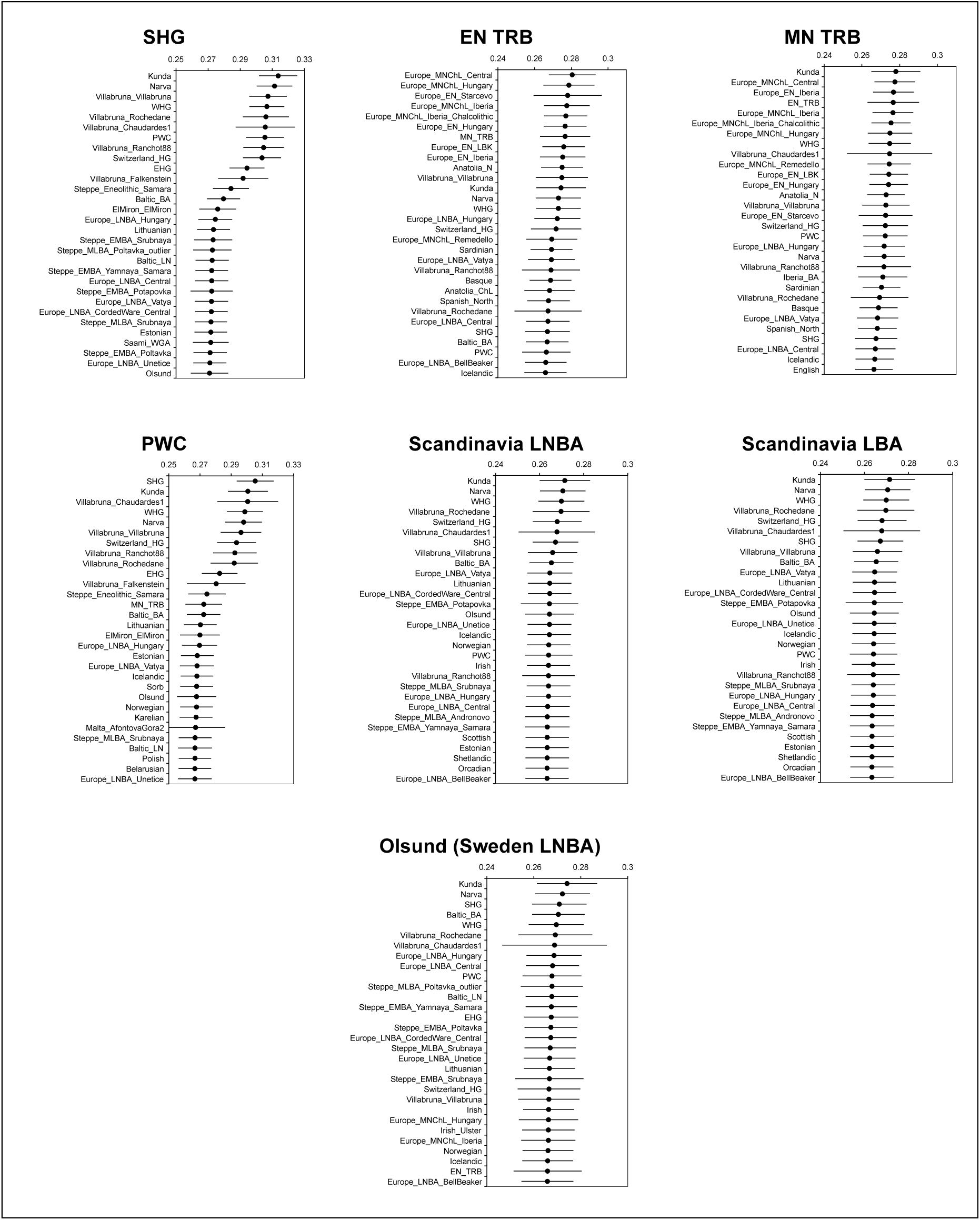
Outgroup *f*_3_-statistics for ancient Scandinavian populations. Values shown for the statistic *f*_3_(Ancient Scandinavian Population, *X*; Mbuti), where *X* is a modern or ancient population. The thirty highest hits are shown and error bars represent three standard errors.

**Extended Data Figure 4.**
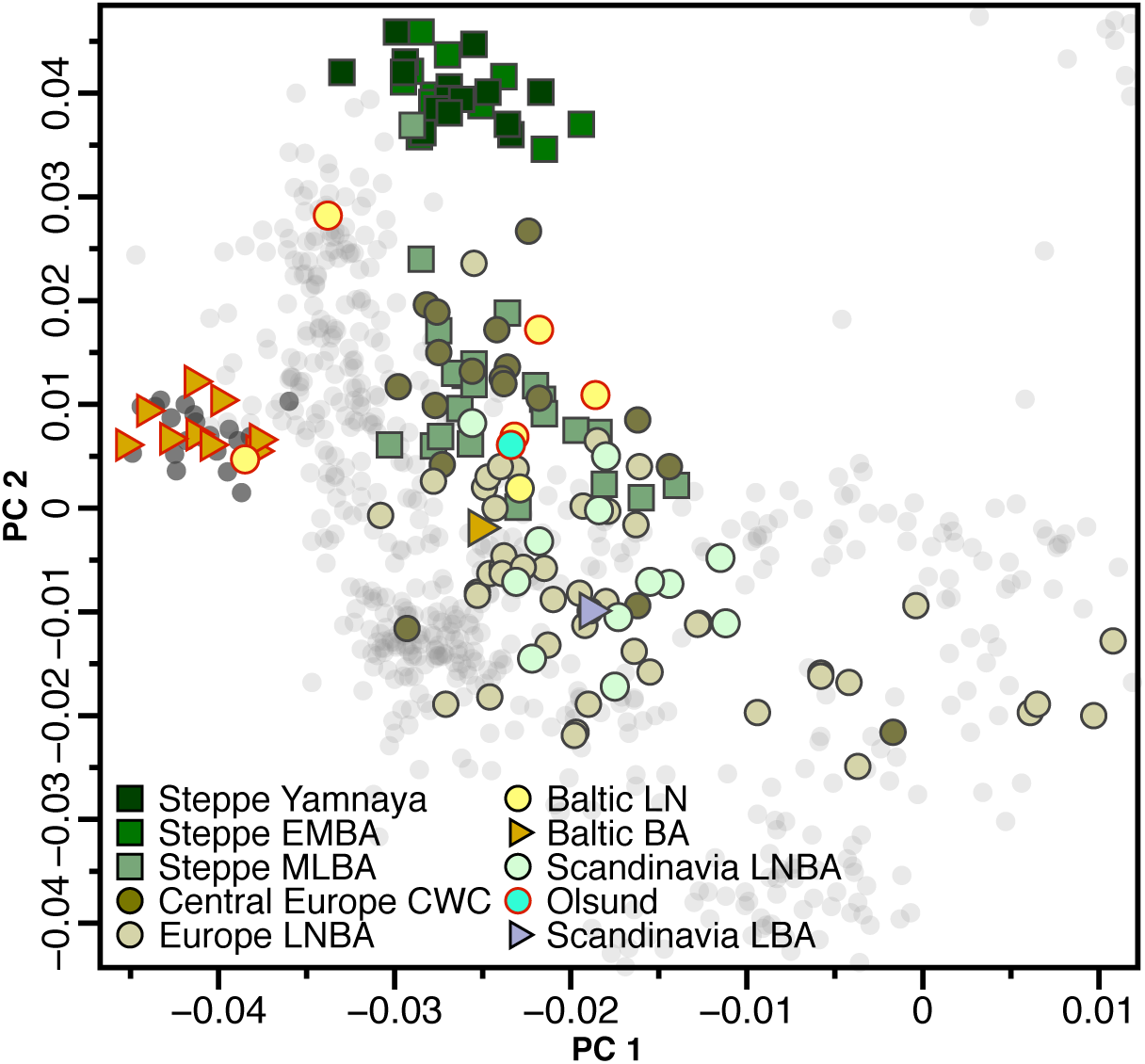
PCA projection of Late Neolithic and Bronze Age Europeans. Zoom-in of corresponding plot in Fig. 2a. Samples introduced in this study are marked with a red outline.

**Extended Data Figure 5.**
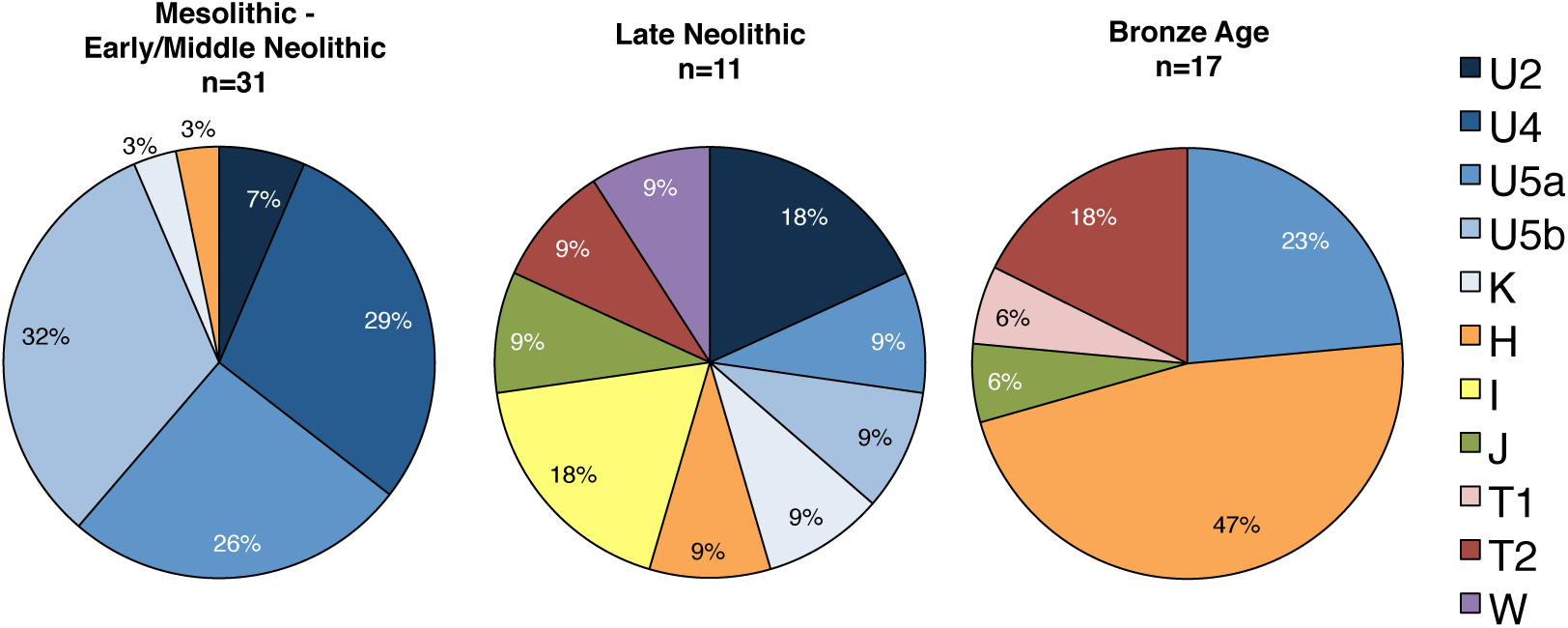
Shift of mtDNA haplogroup frequencies at the onset of the Late Neolithic in the eastern Baltic region. Frequency pie charts of 52 eastern Baltic mtDNA haplogroups generated in this study and seven haplogroups from Bramanti et al. (2009) and Jones et al. (2017).

**Extended Data Table 1.**
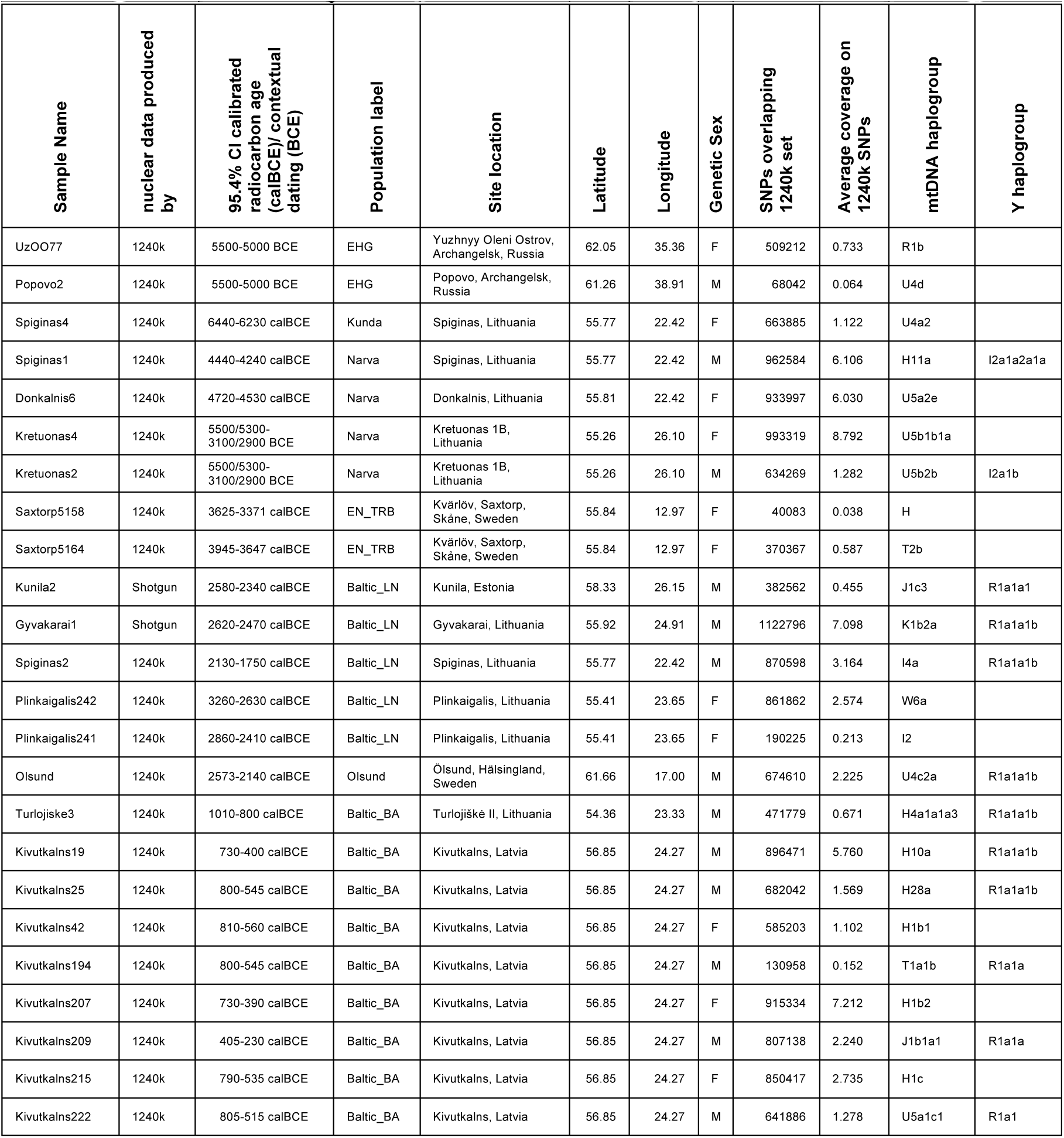
Information on ancient samples for which we report nuclear data in this study.

**Extended Data Table 2.** Allele information of SNPs thought to be affected by selection. Only high-quality (q>30) bases are counted. rs4988235 is responsible for lactase persistence in Europe, the SNPs at *SLC24A5* and *SLC45A2* for light skin pigmentation. The SNP at *EDAR* affects tooth morphology and hair thickness. The SNP at *HERC2* is the primary determinant of light eye color in present-day Europeans. We highlight sites that are likely heterozygous in pink and sites that are likely to be homozygous for the derived allele in blue. DAF= Derived allele frequency.

| Population | Sample | SNP | LCT<br>rs4988235 | SLC45A2<br>rs16891982 | SLC24A5<br>rs1426654 | EDAR<br>rs3827760 | HERC2<br>rs12913832 |
| --- | --- | --- | --- | --- | --- | --- | --- |
|  |  | Ancestral<br>Derived | G<br>A | C<br>G | G<br>A | A<br>G | A<br>G |
| EHG | Uz0077 | Coverage | 3 | 1 | 0 | 7 | 0 |
|  |  | DAF | 0% | 0% | n/a | 0% | n/a |
| Kunda | Spiginas4 | Coverage | 4 | 3 | 2 | 0 | 2 |
|  |  | DAF | 0% | 0% | 0% | n/a | 100% |
| Narva | Spiginas1 | Coverage | 30 | 31 | 6 | 20 | 13 |
|  |  | DAF | 0% | 0% | 50% | 0% | 100% |
|  | Donkalnis6 | Coverage | 34 | 17 | 1 | 21 | 12 |
|  |  | DAF | 0% | 0% | 100% | 0% | 100% |
|  | Kretuonas2 | Coverage | 4 | 5 | 1 | 6 | 2 |
|  |  | DAF | 0% | 100% | 100% | 0% | 100% |
|  | Kretuonas4 | Coverage | 48 | 25 | 4 | 38 | 18 |
|  |  | DAF | 0% | 100% | 75% | 0% | 100% |
| Baltic<br>LNBA | Gyvakarai1 | Coverage | 7 | 5 | 10 | 9 | 8 |
|  |  | DAF | 0% | 60% | 100% | 0% | 100% |
|  | Kunila2 | Coverage | 1 | 0 | 1 | 0 | 1 |
|  |  | DAF | 0% | n/a | 100% | n/a | 0% |
|  | Spiginas1 | Coverage | 30 | 31 | 6 | 20 | 13 |
|  |  | DAF | 0% | 0% | 50% | 0% | 100% |
|  | Plinkaigalis241 | Coverage | 1 | 0 | 0 | 2 | 0 |
|  |  | DAF | 100% | n/a | n/a | 0% | n/a |
|  | Plinkaigalis242 | Coverage | 7 | 5 | 3 | 9 | 2 |
|  |  | DAF | 0% | 0% | 100% | 0% | 0% |
| EN TRB | Kvarlov5164 | Coverage | 1 | 2 | 0 | 0 | 0 |
|  |  | DAF | 0% | 50% | n/a | n/a | n/a |
|  | Kvarlov5158 | Coverage | 5 | 2 | 0 | 1 | 1 |
|  |  | DAF | 0% | 50% | n/a | 0% | 100% |
| Sweden<br>LNBA | Olsund | Coverage | 8 | 4 | 4 | 9 | 2 |
|  |  | DAF | 0% | 0% | 100% | 0% | 0% |
| Baltic<br>Bronze Age | Turlojiske3 | Coverage | 1 | 0 | 0 | 1 | 2 |
|  |  | DAF | 0% | n/a | n/a | 0% | 100% |
|  | Kivutkalns207 | Coverage | 18 | 28 | 4 | 47 | 15 |
|  |  | DAF | 61% | 50% | 100% | 0% | 100% |
|  | Kivutkalns19 | Coverage | 26 | 19 | 5 | 19 | 14 |
|  |  | DAF | 0% | 100% | 100% | 0% | 64% |
|  | Kivutkalns194 | Coverage | 1 | 0 | 0 | 1 | 0 |
|  |  | DAF | 100% | n/a | n/a | 0% | n/a |
|  | Kivutkalns215 | Coverage | 7 | 11 | 2 | 17 | 3 |
|  |  | DAF | 57% | 0% | 100% | 0% | 100% |
|  | Kivutkalns209 | Coverage | 5 | 11 | 2 | 10 | 3 |
|  |  | DAF | 20% | 100% | 100% | 0% | 100% |
|  | Kivutkalns25 | Coverage | 6 | 6 | 1 | 10 | 1 |
|  |  | DAF | 100% | 100% | 100% | 0% | 100% |
|  | Kivutkalns222 | Coverage | 3 | 4 | 0 | 7 | 3 |
|  |  | DAF | 33% | 100% | n/a | 0% | 100% |
|  | Kivutkalns42 | Coverage | 3 | 9 | 0 | 4 | 2 |
|  |  | DAF | 33% | 100% | n/a | 0% | 0% |

